# Temporal, environmental, and biological drivers of the mucosal microbiome in a wild marine fish, *Scomber japonicus*

**DOI:** 10.1101/721555

**Authors:** Jeremiah J Minich, Semar Petrus, Julius D Michael, Todd P Michael, Rob Knight, Eric E. Allen

## Abstract

Changing ocean conditions driven by anthropogenic activity may have a negative impact on fisheries by increasing stress and disease with the mucosal microbiome as a potentially important intermediate role. To understand how environment and host biology drives mucosal microbiomes in a marine fish, we surveyed five body sites (gill, skin, digesta, GI, and pyloric caeca) from 229 Pacific chub mackerel, *Scomber japonicus*, collected across 38 time points spanning one year from the Scripps Institution of Oceanography Pier, making this the largest and longest wild marine fish microbiome survey. Mucosal sites had unique communities significantly different from the surrounding sea water and sediment communities with over 10 times more diversity than sea water alone. Although, external surfaces such as skin and gill were more similar to sea water, digesta was similar to sediment. Both alpha and beta diversity of the skin and gill was explained by environmental and biological factors, especially sea surface temperature, chlorophyll a, and fish age, consistent with an exposure gradient relationship. We verified that seasonal microbial changes were not confounded by migrations of chub mackerel sub-populations by nanopore sequencing a 14 769 bp region of the 16 568 bp mitochondria. A cosmopolitan pathogen, *Photobacterium damselae*, was prevalent across multiple body sites all year, but highest in the skin, GI, and digesta between June and September. Our study evaluates the extent which the environment and host biology drives mucosal microbial ecology, establishing a baseline for long term monitoring surveys for linking environment stressors to mucosal health of wild marine fish.

## Introduction

Pacific chub mackerel, *Scomber japonicus* (Houtuyn 1782), is an economically and ecologically important, cosmopolitan, marine coastal-pelagic fish found in the temperate and tropical waters of the Pacific, Atlantic, and Indian Oceans [1, 2]. *S. japonicus* is currently the fifth largest commercial fishery (purse-seine) in the world [3], processed for human consumption and animal food. Historically in the US, *S. japonicus* was a prominent commercial fishery, but has been on the decline since the 1980s due to a collapse in spawning and fishery stock biomass leading to the last US mackerel canary closing in 1992 [4]. The boom and busts of the fishery have been attributed to large scale environmental factors such as Pacific decadal oscillation, North Pacific gyre oscillation, sea surface temperature, sea level, upwelling, and chlorophyll *a* [5–8]. Juveniles grow quickly reaching 50% of total growth by the first 1.5 years of life with larval growth highest in warmer water (16.8 - 22.1°C) [9]. Larvae eat copepods and zooplankton [3], while juveniles and adults consume primarily small fish and pelagic crustaceans [2]. *S. japonicus* are an important prey item for marine mammals, sea birds, and higher trophic fish such as tunas and sharks [2]. In the eastern North Pacific, *S. japonicus* migrate North in the summer and South in the winter [10] with seasonal offshore migrations occurring from March to May. Climate change and warming oceans likely have contributed to stocks shifting to more northerly migrations [4]. Modeling has shown that nearly 90% of the *S. japonicus* catch was explained by temperature (28-29.4 °C), salinity 33.6-34.2 psu, and chlorophyll *a* of 0.15-0.5 mg/m^3^ [8] whereas survival rates to one year recruits was highly associated with low plankton biomass [11]. *S. japonicus* are ecologically and commercially important while occupying broad environmental gradients. This combined with their relative ease of collection make them an excellent model organism to study the environmental and biological drivers of microbiome diversity in a marine vertebrate.

The primary mucosal environments of fish include the gills, skin, and throughout the gastrointestinal tract, all of which are important to fish health. Disease resistance in the host is promoted in the mucus through continual epithelial shedding and immune cell regulation [12, 13]. The mucus is an important physical barrier to the environment and is generally thought to be colonized with a unique microbiome [14]. The skin and gut both have mucosal associated lymphoid tissues which produce IgT+ B cells protecting the host from invasion of mucosal microbiota [15]. The establishment of microbiomes on mucosal sites is a function of exposure and successful colonization. Mucosal environments within a fish, have varying levels of environmental exposure such as habitat (sea water, sediment, kelp forests), with gills and skin having different exposure rates than the gastrointestinal tract. The GI tract, however will have varying exposure to nutrients during digestion of food. Successful colonization within mucosal sites will further be driven by variables regulated by the host which can include different physiological conditions of the host, thus microbial communities are likely to be reflected. Various protective enzymes related to the innate immune response including lysozymes, proteases, phosphatases, esterases, and sialic acid can be differentially abundant in the mucus depending on the host fish exposure to environmental microbes [16].

To understand the full microbiome potential of a given host, it is important to evaluate the variability longitudinally throughout an entire season (year), and then to continue sampling throughout consistent periods for multiple years. Including long term biological monitoring of commercially and ecologically important marine fish to complement the 100 years of sea water data taken from the SIO pier will be important for understanding marine ecosystem dynamics. Although most ecological studies since 2004 are less than a year and have sampling frequencies of 1 month or greater [17], we have designed our study to include 38 sampling frequencies across 1 year. Previous work looking at seasonal or temporal microbiome changes in the marine environment has focused on free-living pelagic seawater microbes [18]. Gilbert *et al.* found that day length described over 65% of microbial community diversity with richness highest during winter months in the North Atlantic [19]. Very few time series datasets spanning an entire year exist for analyzing the host-associated microbiome. Within humans, most seasonally-active microbes in the gut are associated with populations spending more time outdoors suggesting that seasonal variance in the environment has a greater influence on those with higher environmental exposure [20]. In freshwater fish, lower microbial diversity and altered composition in the gut was associated with warmer summer months in tilapia reared in earthen ponds [21]. In salmon however, no seasonal variations of gut microbiota composition were detected although alpha diversity was highest during warm water months [22]. To date, no systematic analysis of the temporal variability in wild fish microbiomes has been done previously.

The purpose of this study was to quantify the effects of the environmental and biological drivers across five unique mucosal body sites in a marine fish over a longitudinal time course spanning one year. From Jan 28 2017 to Jan 26 2018, 229 pacific chub mackerel, *Scomber japonicus*, were collected off the SIO pier across 38 sampling events. Mucosal microbiome communities were sampled from five body sites including gill, skin, digesta, gastrointestinal tract, and pyloric caeca within each fish. Regional coastal sampling of sea water and marine sediment microbial communities were collected to compare mucosal communities to potential environmental sources. Microbiome processing was performed using the Earth Microbiome protocol using the 16S rRNA gene V4 region. Water conditions (salinity, temperature, pressure, and chlorophyll a) and fish biometrics (length, mass, condition factor, age) were collected and compared to mucosal microbiomes to determine significant ecological drivers. We evaluate both alpha diversity measures (Shannon entropy and Faith’s Phylogenetic diversity) and beta diversity (weighted and unweighted UniFrac) to assess these changes. In addition, we calculate microbial gamma diversity across body sites and time to understand effects of sampling effort on capturing true host-microbiome diversity. Our results show that mucosal communities across body sites are highly differentiated in a single species of fish and that seasonal environmental drivers partially account for this differentiation.

## Materials and Methods

### Sample collection *S. japonicus* time series

From Jan 28 2017 to Jan 26 2018, 1-8 *Scomber japonicus* specimens were collected across 38 sampling events from the end of the SIO Pier (32.867, −117.257). Sea surface water samples were collected from each sampling event and immediately stored on dry ice. Environmental conditions at time of sampling, including sea water temperature, salinity, pressure, and chlorophyll *a* concentration, were collected using publicly available data from the Southern California Coastal Ocean Observing System (SCCOOS) SIO pier shore station data archive (http://www.sccoos.org) (Fig 1a). Fishing occurred at or near sunset with exact times recorded in the metadata (see Qiita Study ID 11721 for full metadata). Fish were caught using hook and line with a Sabiki rig, immediately euthanized upon landing using accepted protocols according to AVMA guidelines and stored on dry ice. Individual fish were wrapped in aluminum foil and handled with gloves prior to storage on dry ice to minimize contamination and then stored long term at −80°C for up to 6 months prior to dissection. Upon processing, frozen fish were weighed and measured, along with calculation of Fulton’s condition factor, which is a proxy for fish health [23, 24]. Age was estimated using fish length as derived from the most recent Pacific chub mackerel stock assessment [4] (Fig 1b-c) [25] where otoliths were compared to 25 fish individuals per catch across multiple years (1962-2008). Specifically, the von-Bertalanffy equation was used with two separate growth coefficients: LA = L∞ (1 − e −k(A-to)) where LA=length at age, L∞=theoretical maximum length of fish, k = growth coefficient, to = theoretical age when length is 0 mm. After 30 minutes of thawing the fish, a cotton swab (Puritan, Cat #806-wc) was swiped back and forth five times along the left gill and then put directly into a 2 ml Mo Bio PowerSoil (Mo Bio, Cat # 12888) bead beating tube. The skin was also swabbed in a 3 cm × 3 cm area on the left side behind the gill and above the pectoral fin (Fig 1d). After carefully dissecting the fish with a new razor blade, the last 3 cm of GI tract was cleared and the digesta sampled. This same distal portion of GI tract was cut and also sampled. Lastly, an approximate 50 mg sample of pyloric caeca was sampled from the fish and placed in a tube. The tubes were then stored at −80°C until DNA extraction commenced. For additional environmental controls, surface seawater and sediment samples were collected across two time points (Dec 8 2017 and Jan 12 2018) at 30 coastal locations, approximately 200 m offshore, spanning 10 km throughout San Diego including soft bottom, reef, river mouth, and bay areas.

**Figure 1.**
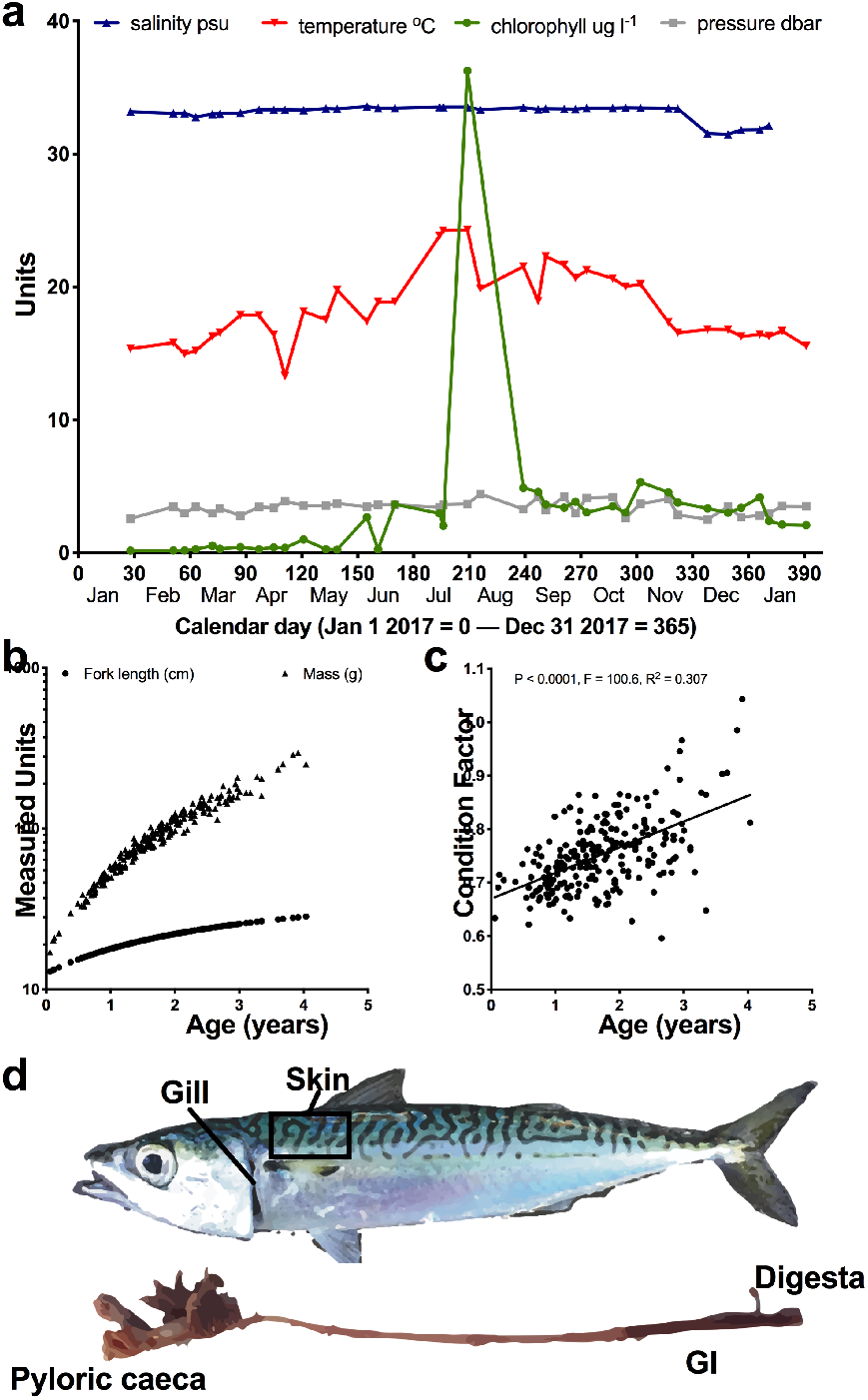
Environmental sampling design. Throughout 2017 and into early 2018, 229 Pacific chub mackerel, *S. japonicus*, were caught across 38 sampling events from the SIO pier. (a) Pier sea water measurements of temperature, salinity, pressure, and chlorophyll a were collected using the scoos.org database. (b) Ages of *S. japonicus* were inferred from fish lengths. (c) Condition factor was calculated for each fish based on length and mass. (d) Mucosal microbiome samples were collected from five body sites including gill, skin, pyloric caeca, GI biopsy, and fecal or digesta material removed from the lower GI.

### Microbiome processing

Samples were processed using the standard 16S rRNA gene Earth Microbiome Protocol (EMP), with only slight modifications (http://www.earthmicrobiome.org). Specifically, genomic DNA was extracted using a hybrid approach where lysis is performed in 2 ml bead beating tubes and then cleanup performed using the KingFisher robot to reduce well to well contamination (Minich et al, in prep). The initial cell lysis steps were performed in single-tube reactions (instead of 96-well plate format) followed by transfer to plates for the standard magnetic bead cleanup on the KingFisher robots using the Mo Bio PowerMag kit (Mo Bio, Cat # 27000-4-KF) which has improved limits of detection for low biomass samples [26]. The EMP extraction procedure includes modifications including the use of RNaseA during lysis and a 10 minute incubation at 65°C prior to bead beating. All sample batches had positive and negative controls included with each extraction set so that sample exclusions based on read counts could be calculated [26]. Extracted gDNA was then PCR amplified using the EMP 16S V4 515f/806rB bar-coded primers [27, 28]. The miniaturized PCR method, which generates libraries at a 58% cost reduction of $1.42 per sample, was used for all samples that included the use of the Echo-550 instrument to do triplicate 5 ul PCR reactions [29]. Amplicons were quantified using a pico green assay, and then 2 ul of each sample was equally pooled into a single tube. This final pool was then cleaned up to remove dNTPs and primer dimers using the QIAquick PCR purification kit (Qiagen, Cat# 28106). Final pools contained up to 768 samples which were then sequenced on an Illumina MiSeq using a 2×150 bp strategy (300-cycles v2 kit, Illumina, San Diego, CA). Bioinformatic processing of samples occurred using Qiita [30] and QIIME 1.9.1 or QIIME 2.0 [31], with the first 150 bp read trimmed to 150 bp and processed through deblur [32], a *de novo* sOTU picking method. A phylogenetic tree of the 16S sOTU single-sequence tags was created using SEPP (SATé-Enabled Phylogenetic Placement) [33]. Rarefaction levels were empirically determined by calculating the read counts at which 90% of the reads from the DNA extraction positive controls map back to the positive controls [26].

### Summary microbiome statistics

Alpha, beta, and gamma diversity of microbial communities was measured [34]. Alpha diversity was calculated using measures of Shannon [35] and Faith’s Phylogenetic Diversity [36] while beta diversity was calculated using weighted and unweighted UniFrac [37, 38] distance and visualized in Emperor [39]. Alpha and beta diversity statistical significance was tested using Kruskal-Wallis test [40]. Taxonomies were classified in Silva [41] using the Greengenes and RDP databases [42] using the following parameters: minimum identity with query sequence (0.95), number of neighbors per query sequence (10), greengenes-reference NR database, search kmer-candidates (1000), lca-quorum (0.8), search-kmer-length (10), search-kmer-mm (0), search-no-fast, reject sequences below 70%.

### Statistical analysis of environmental and biological drivers of fish mucosal microbiomes

To evaluate the extent to which the environment and biology of the fish influences the microbial communities of the various body sites, both alpha diversity and beta diversity were analyzed. Only samples which had environmental values for all water conditions (temperature, salinity, pressure, and chlorophyll a) and biological conditions (weight, fork length, condition factor, and age) were included. Thus, some samples had to be excluded due to temporary failure of the chlorophyll *a* fluorometer instrument on the SIO pier (August 4 2017). Alpha diversity measures for each body site were independently verified and tested to ensure they met the assumptions for the General Linear Model (GLM). Specifically, to test for normally distributed residuals, sets were analyzed using the R package Library(car) [43] and run through the Shapiro-Wilk Normality test [44]. To evaluate and test for homoscedasticity, the non-constant error variance test (ncvTest) commonly known as the Breusch-Pagan test was used [45]. To meet GLM criteria, the Faith’s Phylogenetic Diversity samples were log-transformed. Both Shannon and Faith’s PD were then processed through the GLM in R while controlling for collinearity of variables. Individual R^2^ values and P-values for each environmental and biological variable are reported along with total R^2^, F-statistic, and P-values for all variables. Gill samples using Shannon diversity and GI samples using Faith’s PD were excluded from analysis due to not meeting required assumptions of the GLM. To evaluate the effects of environmental and biological variables on beta diversity, we assessed both unweighted and weighted UniFrac for each body site independently using Adonis, a non-parametric analysis of variation method, [46] in QIIME 1.9.1 [31] and Calypso [47].

### Validation of *Photobacterium damselae* sOTU phylogeny

To validate the taxonomy assignments of five *Photobacterium* sOTUs in our dataset, we performed multiple sequence alignment (Neighbor-Joining) of a 148 bp region of the 16S rRNA gene v4 region from eight other strains. Specifically, we used default settings (nucleotide scoring 200 PAM / k=2, Gap opening penalty 1.53, offset value = 0, ‘nzero’ where Ns have no effect on alignment score) in the MAFFT alignment tool [48, 49]. The phylogenetic tree was visualized using Phylo.io [50]. The comparison bacteria strains included: two pathogenic *Photobacterium spp.* isolates (*P. damselae* ATCC 33539T, Genbank X74700.1; *P. damselae*, Genbank D25308.1), four non-pathogenic *Photobacterium spp*. (*P. leiognathi* strain ATCC 25521, Genbank NR_115541.1; *P. angustum* ATCC 25915T, Genbank X74685.1; *P. phosphoreum* strain ATCC 11040, Genbank NR_115205.1; *P. rosenbergii* strain CC1, Genbank NR_042343.1) and two outgroup *Vibrio* species (*Vibrio pelagius* strain ATCC 25916, Genbank NR_119059.1; *Aliivibrio fischeri* strain ATCC 7744, Genbank NR_115204.1) which were identified from various studies [51–53].

### Population genetics of *S. japonicus*

We developed a high-throughput two fragment, mitochondrial amplicon workflow for the Oxford Nanopore long read sequencer. A total of 96 gDNA skin mucus samples, spanning 5 collection months (Aug 27 2017 – Jan 26 2018) were amplified in 192 separate 10 ul PCR reactions using 1 ul gDNA, 5 ul NEB Long Amp mastermix (NEB, Cat# M0287S), and 3.4 ul molecular grade water and one of two primer combinations. The first mtDNA fragment (96 PCR reactions) used 0.4 ul of 10uM forward primer (SJ_F1_655: 5’-TTT CTG TTG GTG CTG ATA TTG | CAA ACC TCA CCC TCC CTT GTT-3’) and 0.4 ul of 10 uM reverse primer (SJ_R1_7653: 5’-ACT TGC CTG TCG CTC TAT CTT | CAC CAC TAT TCG GTG GTC TGC-3’). The second fragment (96 PCR reactions) used 0.4 ul of 10uM forward primer (SJ_F2_7425: 5’-TTT CTG TTG GTG CTG ATA TTG | CTC CCT GCC GTC ATT CTT ATC) and 0.4 ul of 10 uM reverse primer (SJ_R2_15424: 5’-ACT TGC CTG TCG CTC TAT CTT | CGA CGA CTA CGT CTG CGA CAA). All primers have ONT adaptor regions on the first 27 bases as indicated by ‘|’. All PCR reactions followed the following protocol: 94 °C 3 minutes, 25 cycles of 94 °C 30s, 60 °C 30s, 65 °C 8:20, a final extension of 68 °C 10 minutes followed by storage at 4 °C. Following the first PCR, a second 5 ul PCR reaction was conducted for each of the 96 samples by combining 2.5 ul NEB Long Amp mastermix, 0.1 ul of a unique barcode (Oxford Nanopore PCR Barcode kit 01-96, batch DK601001 brown box) and finally pooling 1.2 ul of each PCR product (first + second fragment). Barcodes were transferred to the PCR reaction plate using the acoustic liquid handler Labcyte Echo 550. The same PCR reaction was used. A final 2 ul of sample was pulled from all samples and processed through the QiaQuick PCR purification kit (Qiagen, Cat#28106) and run on a 1% agarose gel to confirm size. The pool was then run on a used MinION using the 1D PCR barcoding protocol (SQK-LSK109). Samples were demultiplexed, uploaded to Galaxy [54], and aligned against the *Scomber japonicus* reference mitochondria genome (NC_013723) using LASTZ aligner (Galaxy version 1.3.2)[55] using defaults. Consensus sequences were visualized, calculated, and exported using the quick consensus mode in Integrated Genome Viewer [56]. Samples with either less than an average of 10x coverage or samples with more than 20 ambiguous basepairs (Ns) were excluded from the analysis (n=2, BC52, BC82). A phylogenetic tree of all 91 *S. japonicus* samples along with three reference *S. japonicus* mitochondrial genomes from NCBI (AB488405.1, NC_013723.1, AB102724.1), and two outgroup species *Scomber colias* (NC_013724.1) and *Scomber australasicus* (AB102725.1) was generated using MAFFT [49] (NJ conserved sites 12388, Jukes-Cantor substitution model, bootstrap 100) and visualized with Phylo.io using default parameters [50].

All microbiome data is publicly available through Qiita (sample ID 11721, prep ID 4638), EBI (ERP109537), and NCBI (BioProject PRJEB27458).

## Results

### Microbial diversity associated with a marine pelagic fish across body sites over one year

From January 2017 to January 2018, 229 wild *Scomber japonicus* were collected from the SIO pier across 38 sampling events at approximately five fish per week, although actual takes varied due to weather and other constraints. Sea water temperature, salinity, pressure, and chlorophyll *a* were recorded using the SCOOS online database (Figure 1a). Fork length and mass were recorded and approximate age of the fish determined from length (Figure 1b). The condition factor of the fish was positively associated with older fish (P<0.0001, R^2^=0.307) (Figure 1c). Along with paired sea water samples, mucosal microbiome samples were sequenced from the gill, skin, digesta, GI, and pyloric caeca of each fish (Figure 1d).

A total of 612 samples resulting in 18 857 sOTUs, processed with the miniaturized PCR method, passed the sample exclusion criteria. Sample exclusion criteria was based on the KatharoSeq method where the read counts from DNA extraction positive controls of varying cell counts was compared to compositional read out and the read count at which 90% of the reads mapped appropriately was chosen as the rarefaction depth, which was 1 362 reads (Supplementary Figure 1). Alpha diversity measured by Faith’s PD, was significantly different when compared across mackerel body sites and sea water (Kruskal-Wallis, P<0.0001, KW statistic 87.48) (Figure 2a). Gill, skin, and digesta samples had higher diversity than the GI and pyloric caeca samples while gill and digesta had higher diversity than sea water (Figure 2a). Beta diversity indicates the gill and skin mucosal samples were most similar to sea water while digesta, GI, and pyloric caeca were uniquely clustered and also had higher within body site variability (Figure 2b). Some of the digesta samples also appeared to cluster more closely to sediment samples. When tested, skin followed by gill samples were most similar to sea water samples whereas the digesta samples were most similar to sediment (Figure 2c). To understand sample size requirements for capturing novel microbial diversity in fish, we compared accumulation of microbial richness over the one-year sampling period across all sample types. Overall microbial richness in the gill, skin, GI, pyloric caeca, and sea water appeared to level off after only a couple months (20-50 samples) whereas digesta samples continued to increase perhaps requiring another few years of data collection to approach saturation. For comparison, we included gill, skin, and digesta samples from 14 other local San Diego species of fish (right of the dotted line) (Figure 2d). While digesta diversity increased with the addition of the first new species it followed a similar trend while gill and skin samples did not increase much suggesting an overall conservation of microbes in other species of fish. Lastly, total Gamma diversity or richness was calculated for all samples in this study showing that sediment samples had the most microbial diversity followed by mackerel digesta and mackerel gill. The total unique microbial diversity in a single species of fish, *S. japonicus*, was 8.8 fold more than sea water (9 172 vs. 1 039 sOTUs) (Figure 2d), demonstrating the potential for microbial discovery within and upon fish hosts in the ocean.

**Figure 2.**
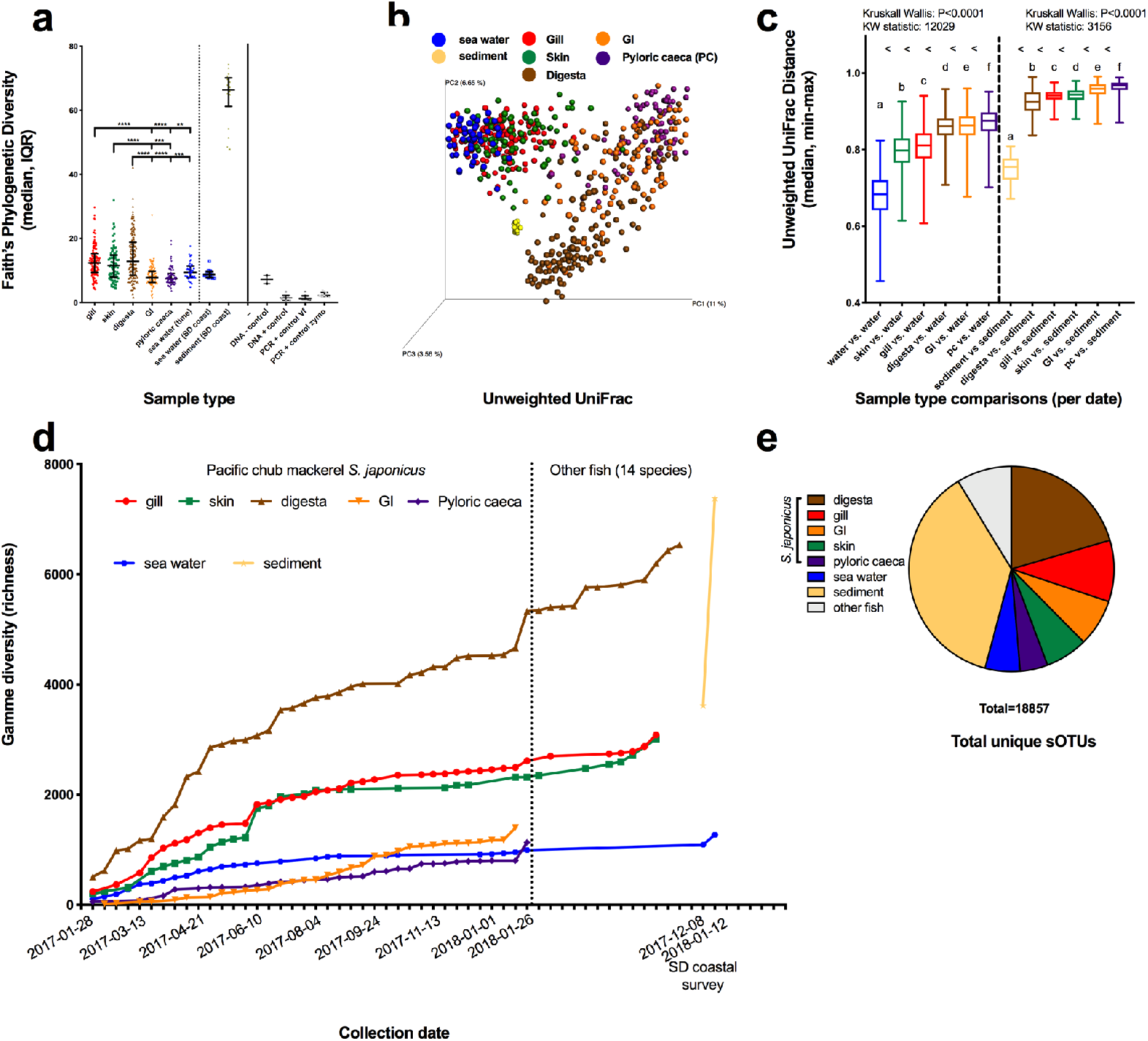
Microbial diversity of coastal environmental controls and *S. japonicus* mucosal microbiome. (a) Alpha diversity was calculated using Faith’s Phylogenetic diversity metric in Qiime 1.9.1 with the median and interquartile range displayed. (b) PCoA plot of beta diversity as calculated using Unweighted UniFrac with a rooted phylogenetic tree inserted using the SEPP method in Qiita and Qiime1.9.1. (c) Distances of mucosal microbial communities (gill, skin, digesta, GI, and pyloric caeca) compared to sea water and sediment samples using non-parametric Kruskal-Wallis test. (d) Accumulation of total microbial diversity across chronological sampling events within fish (*S. japonicus* and 14 other species) mucosal sites, water samples, and sediment. (e) Proportion of unique microbial diversity (sOTUs) contributed by body site or environment to the whole dataset.

### Environmental and biological drivers of the *S. japonicus* mucosal community

We next quantified the combined and specific effects of four environmental variables including chlorophyll *a* concentration, sea water temperature, salinity, and pressure along with four biological variables including fish age, fork-length, mass, and condition factor on the fish-associated mucosal microbiomes. Alpha diversity measures were assessed using the General Linear Model (GLM). For alpha diversity measures of Shannon diversity, skin mucus was significantly influenced by the factors (P<0.001, R^2^ = 0.38, F-stat 6.595), with chlorophyll *a* having a negative association and temperature a positive association (P<0.0001, P=0.0004), respectively. Gill samples were not assessed (grey line Table 1) because the Shannon diversity did not meet the assumptions of normally distributed residuals (Shapiro test P <0.05) and was not homoscedastic (Breusch-Pagan P<0.05) (Table 1). For the alpha diversity measure of Faith’s phylogenetic diversity (PD), which takes into account phylogenetic diversity with richness, all data was log transformed to meet the assumptions of the GLM. The gastrointestinal tract samples however, still did not meet the assumptions as the residuals were not normally distributed (Shapiro test P<0.05), thus were excluded from analysis (grey line Table 1). Both gill, skin, and pyloric caeca Faith’s PD were significantly influenced by the measured factors (gill: P<0.0001, R^2^=0.33, F-state=7.042; skin: P=0.00039, R^2^=0.26, F-stat=4.239; pyloric caeca: P=0.00891, R^2^=0.22, F-stat=2.972). The gill sample diversity was negatively associated with Chlorophyll a concentration (P=0.00549). Skin was negatively associated with Chlorophyll *a* concentration (P=0.00182) and age (P=0.00811), while positively associated with temperature (P=0.04434). The pyloric caeca was positively associated with age (P=0.04787) and temperature (P=0.00305) while negatively associated with salinity (P=0.04921).

**Table 1.**
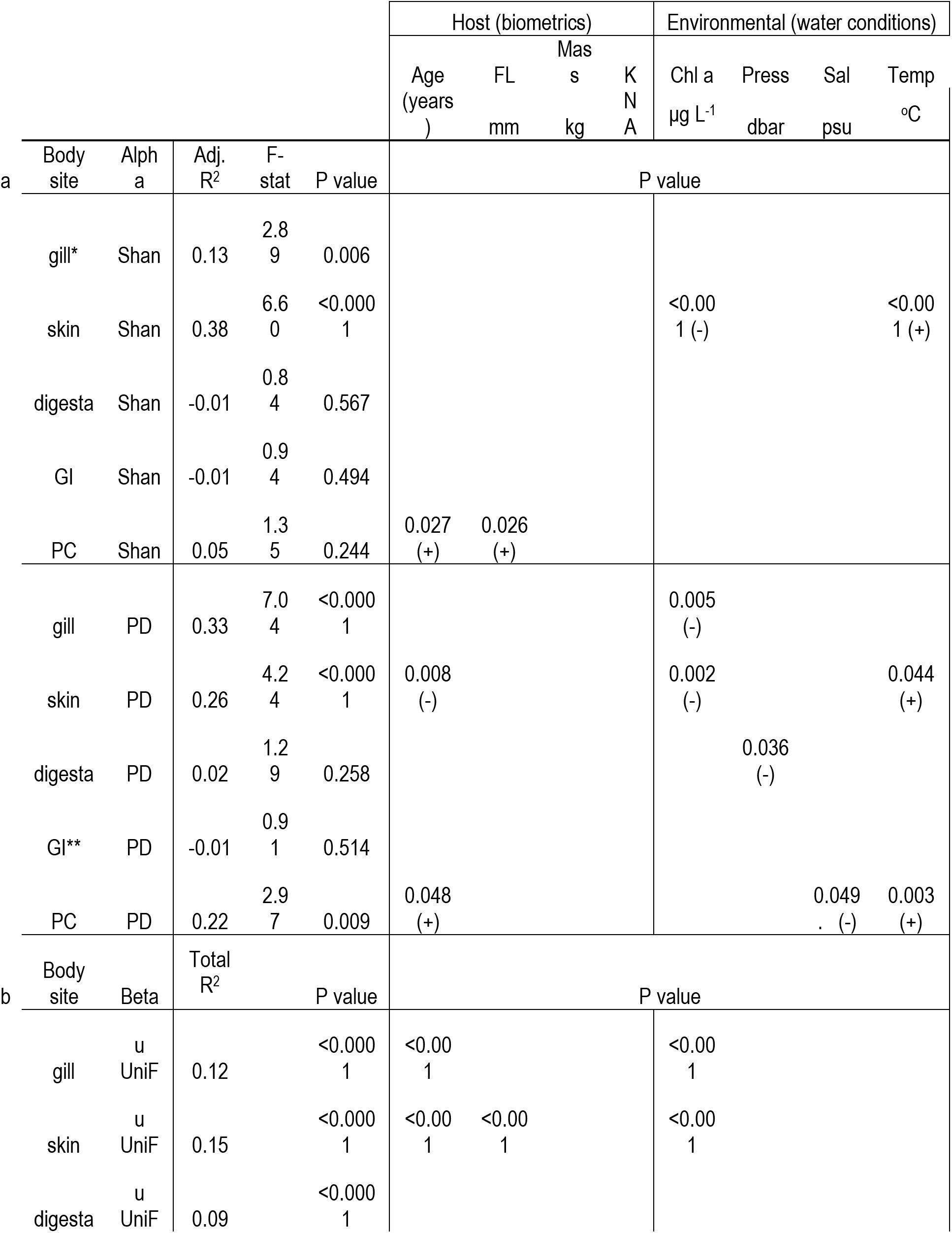
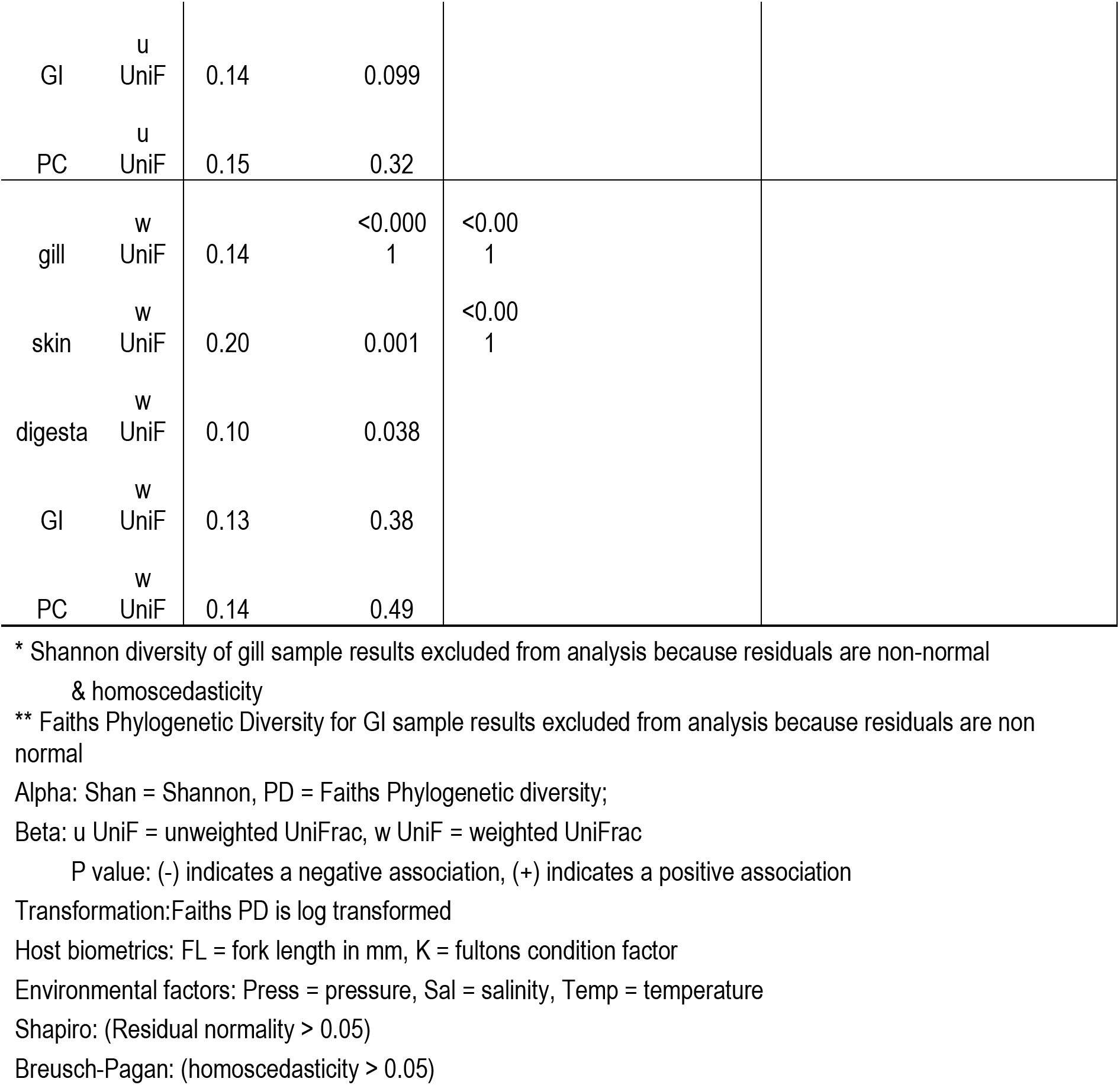
Quantification of environmental and biological variables on fish mucosal microbiomes as measured by (a) alpha diversity with Generalized Linear Model and (b) beta diversity with multivariate statistics (Adonis).

The extent to which environmental and biological variables explain microbial diversity was also assessed for Beta diversity including both unweighted UniFrac and weighted UniFrac distances. The Adonis permutational multivariate statistical analysis was used to test overall significance along with variance explanation by factor. Unweighted UniFrac distance measures showed that gill, skin, and digesta samples were influenced by measured factors (Adonis, P<0.0001, R^2^=0.12, R^2^=0.15, R^2^=0.09). The gill was primarily driven by Chlorophyll *a* concentration and age while skin was influenced mostly by Chlorophyll *a*, age, and fork-length. For weighted UniFrac distances, both gill (P<0.0001, R^2^=0.14) and skin (P=0.001, R^2^=0.20) were significantly influenced by factors with age being the most significant driver. In summary, the skin mucosal microbiome was significantly influenced by environmental and biological factors in each of the four measures across alpha and beta diversity while gill was significant in each of the three measures. The environmental variables of Chlorophyll *a* followed by temperature had the most frequent influences on microbial communities across body sites while age was the most frequent biological factor (Table 1).

### Population structure of *S. japonicus*

*S. japonicus* are thought to have three spawning populations along the Pacific coast of North America[4], which suggest that our environmental and biological associations could be explained in part by population dynamics over the year. To estimate the changes in *S. japonicus* population structure over our time course we sequenced two fragments of mitochondrial DNA directly from skin mucus gDNA for a total length of 14 769 bp for 93 fish samples landed between Aug 27 2017 and Jan 26 2018. Two samples were removed from the analysis due to having lower coverage (less than 10x) or more than 20 Ns in the consensus sequence (Supplemental Figure 2a). The majority of samples (93%) had at least 100x coverage of the mitochondria target region (Supplemental Figure 2b). Based on near full-length mitochondria data, no population structure was observed, consistent with our sampling of one population of *S. japonicus* over the course of the study (Supp Figure 3).

### Candidate pathogen and probiotic associations

Microbial diversity was unique within the various *S. japonicus* body sites and environment (sea water and sediment) with the top 25 most abundant genera comprising the majority of reads (Figure 3). *Rhodobacteraceae* were found in all environments particularly the seawater with lower levels in the gill, skin, digesta, and GI body sites. The gill was dominated by several sOTUs within the *Shewanella* genera along with microbes from the *Rickettsiales* and *Polynucleobacter* genera. Sea water microbes including *Rhodobacter* and *Pelagibacter* were also present on the gill in lower numbers. Various sOTUs of *Photobacterium* were highly abundant across the skin, digesta, GI, and pyloric caeca. *Enterovibrio*, another fish pathogen, was abundant on the fish skin and pyloric caeca. Digesta samples were comprised of many seawater dwelling Cyanobacteria including *Synecochoccus* but was also high in the family *Pirellulaceae* (Planctomycetes) and sporadically high amounts of the genus *Clostridium*. *Mycoplasma* (Tenericutes) was a dominant genus in the GI and pyloric caeca. Various *Bacillus spp.* and *Lactobacillus spp*. were found to be present across multiple body sites (Supplemental Figure 4). As expected, seawater contained many common groups including *Synechococcus*, *Rhodobacteraceae*, *Pelagibacter*, and *Flavobacteriaceae* while the sediment had consistently higher levels of *Pirellulaceae* (Figure 4).

**Figure 3.**
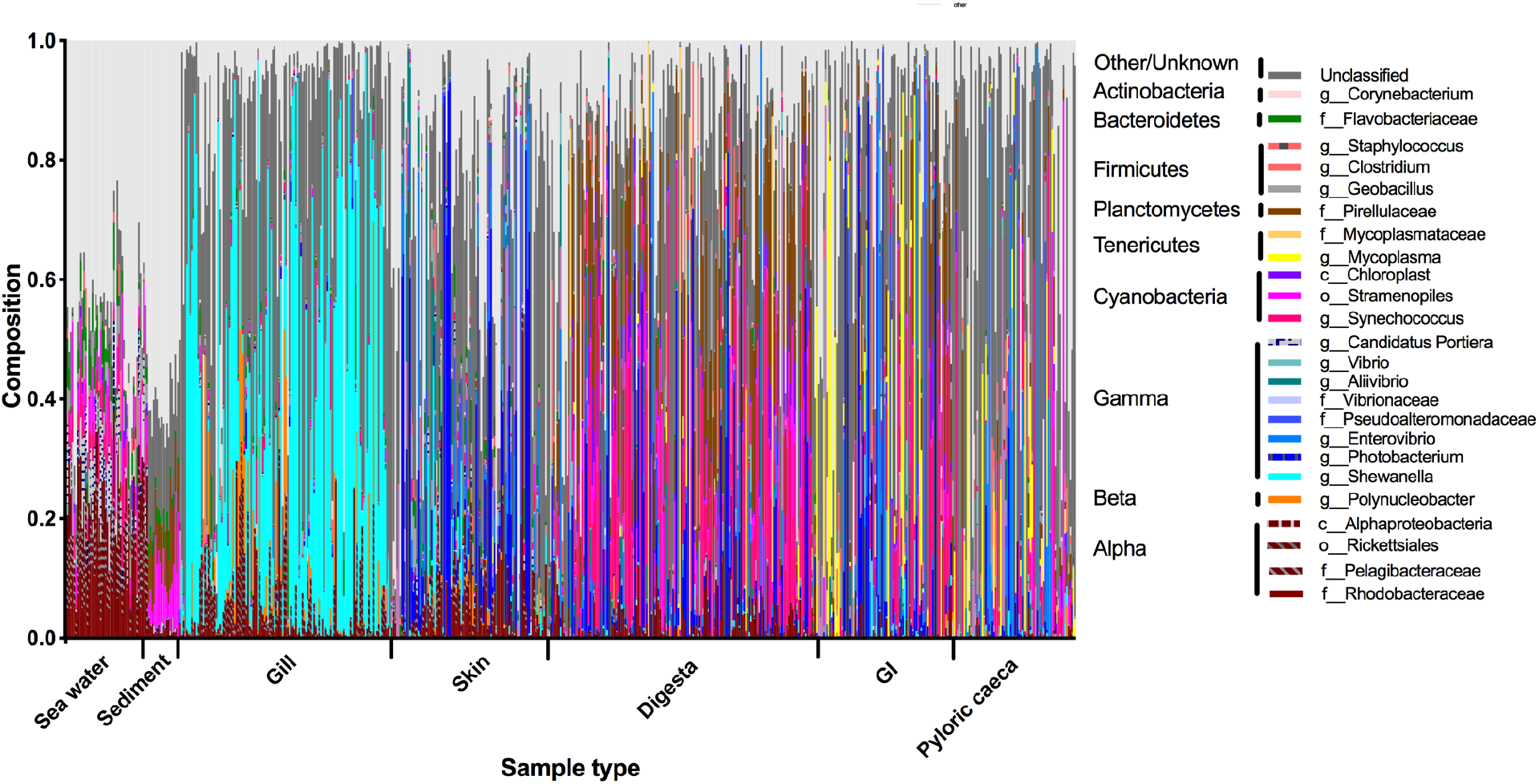
Top 25 most abundant genera across body sites.

**Figure 4.**
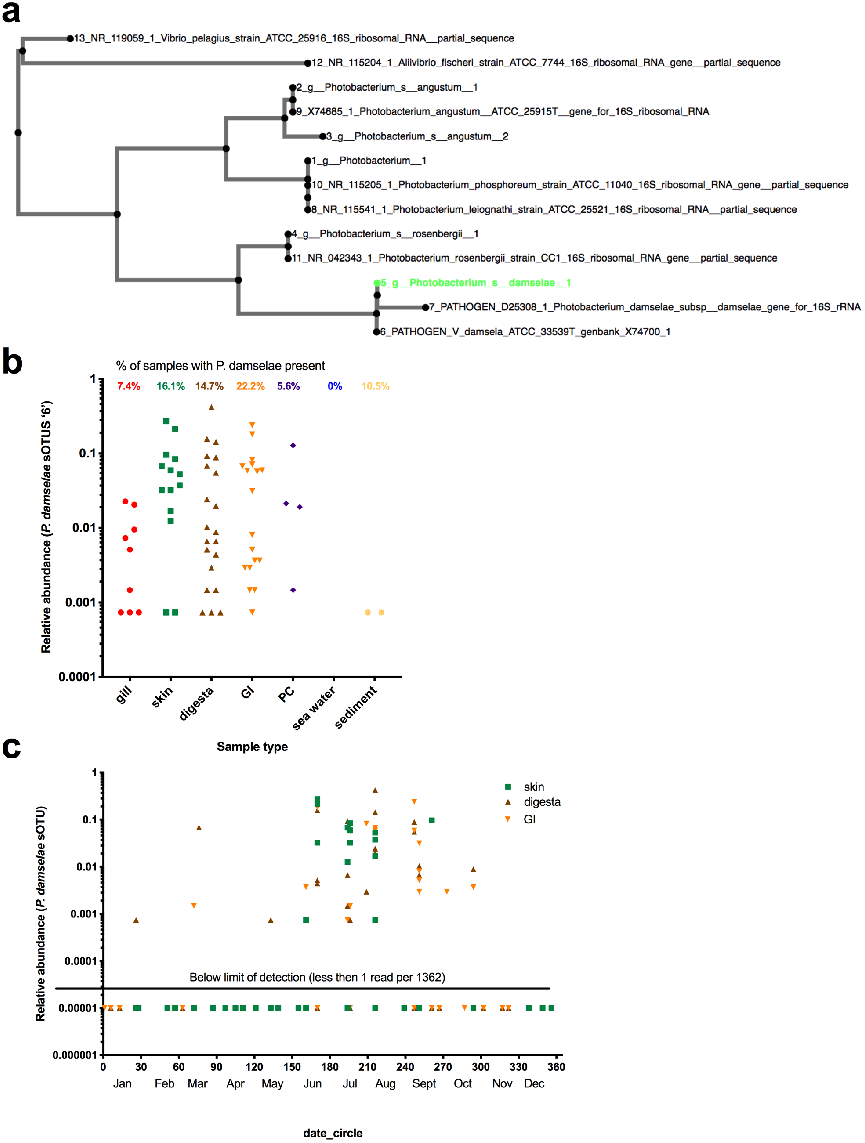
Prevalence of marine vertebrate pathogen, *Photobacterium damselae* on *S. japonicus* body sites throughout the sampling effort. (a) Validation of *P. damselae* 148 bp v4 region by way of phylogenetic comparison to two known pathogen isolates and and non-pathogenic strains. Total relative abundance of *P. damselae* sOTU across five body sites and environments for successfully sequenced samples. Total prevalence or percent of samples with *P. damelsae* present is also calculated for each sample type and displayed at top of graph (7.4% gill samples, 16.1% skin, 14.7% digesta, 22.2% GI, 5.6% pyloric caeca, 0% water, 10.5% sediment). (c) Proportion of microbial community comprised of *P. damselae* sOTU across the most prevalent body sites (skin, digesta, GI) over the sampling effort of 1 year. Relative abundance is calculated as number of *P. damselae* sOTU reads divided by 1362, the rarefraction number. Any samples with 0 *P. damselae* reads are considered under the detection limit and are displayed as equal to 0.00001 relative abundance in order to visualize on the log scale.

There were five highly abundant *Photobacterium* sOTUs in the dataset which prompted further phylogenetic evaluation to elucidate species level assignments. The full 16S rRNA genes of two pathogenic isolates of *P. damselae*, four other Photobacterium species including *P. angustum*, *P. phosphoreum*, *P. leiognathi*, and *P. rosenbergii* and two vibrio species as outgroups were aligned with the five *Photobacterium* unique 150 bp sOTUs. The phylogenetic tree (Figure 4a) of the five Photobacterium sOTUs in this dataset with the known *Photobacterium* strains is able to identify and resolved the taxonomic assignments. The *Photobacterium damselae* sOTU identified in our dataset was 100% identical to the v4 region of 16S rRNA from the two pathogenic strains while distinct from the other *Photobacterium spp*. This *P. damselae* sOTU was identified across various body sites of fish, but was most prevalent in the GI, skin, and digesta samples (present across 22.2%, 16.1%, and 14.7% of samples, respectively) (Figure 4b). For the GI, skin, and digesta samples which had *P. damselae* present, the single *P. damselae* sOTU made up 5.88%, 6.99%, and 5.32% of the total microbial composition, respectively. Further, the temporal enrichment and prevalence of this *P. damselae* sOTU was highest between June and September.

## Discussion

Our study evaluated how the mucosal microbial community of a wild marine fish species is influenced according to environmental and biological variance as experienced over the course of an annual season in coastal temperate waters. Body sites had unique microbial signatures that were uniquely influenced by environmental and biological measures. Alpha diversity was highest in the gill, skin, and digesta communities as compared to the gastrointestinal tract and pyloric caeca. Beta diversity measures demonstrated that fish mucosal sites were primarily driven by body site location and were further unique to the surrounding environment. An exposure gradient was observed with skin and gill surfaces being more similar to the water column while the digesta community more similar to the sediment. Further, the environmental and biological variables best explained variation in the skin and gill microbiomes as opposed to the internal body sites (digesta, GI, pyloric caeca). Lastly, an important fish pathogen, *Photobacterium damselae* was observed in high prevalence on GI, skin, and digesta communities and was associated with the summer months which exhibit higher temperatures and low nutrients. This is the first comprehensive microbiome study of a marine fish that evaluates multiple body sites from a large sample size over an extended time series.

Regardless of environmental conditions, the mackerel mucosal body site was the strongest driver of microbiome diversification, with each site associated with a specific gradient of environmental exposure. The gill and skin communities were most similar to the seawater whereas the gastrointestinal samples were more divergent. This environmental gradient which distinguishes host-associated gut microbes from free-living microbes, was first described in mammalian vertebrates [57]. Environmental exposure gradients have also been shown to influence gut or skin microbiomes in amphibians, fish [58, 59], and other vertebrates [60, 61], whereas our study is the first comprehensive community assessment that explicitly tests this comparison with multiple mucosal body sites in fish. Marine fish differ from other vertebrates in that their microbial exposure rates are greatly elevated compared to terrestrial or freshwater species. Seawater can harbor as many as 1 million cells per ml [62], while coastal sediments can be two-orders of magnitude higher at 100 million cells per cm^3^ [63]. Gill microbial communities may be supported physically by complex morphological structure of laminae and filaments and chemically through gas exchange, ion transport, and waste excretion. Age, phylogeny, diet, and individual have been implicated as influencing the gill microbial community in tropical fish, with *Shewanella* taxa being dominant [64]. Skin microbiomes of marine tropical fish have been also shown to be driven by phylogeny and diet [65]. Digesta and GI samples in *S. japonicus* were the most variable suggesting that either niche differentiation is more static in the gill and skin environments or that microbial turnover is lower. Few studies, however, have evaluated these body sites in temperate marine fish. Discovery of novel microbial lineages and metabolic activity should focus on fish mucosal associated environments, specifically the gill, skin, and digesta communities that had the highest levels of phylogenetic diversity in our dataset. Sediment samples had the highest diversity, yet were most similar, thus having the lowest inter-individual variability.

The environmental and biological variables most explained the skin and gill microbiomes as compared to the internal GI communities. Within the environmental variables, Chlorophyll *a* concentration followed by temperature and salinity were the strongest drivers while age was the most pronounced of the biological metrics. Chlorophyll *a* concentration is a general indicator of primary production and microbial growth or proliferation in the water column. As phytoplankton blooms occur in the ocean due to nutrient enrichments through upwelling, bacterial communities in the water column also change thus altering exposure to fish and other marine animals. While many studies have examined the effects of harmful algal blooms on marine organisms [66], few have quantified the extent of these exposures in the wild. Temperature has been shown to influence marine macroalgae [67] and oyster hemolymph microbiomes [68]. Salinity was one of the first major abiotic conditions shown to drive microbiomes in free-living freshwater versus marine environments [69] and has also been shown to influence fish microbiomes deterministically [70]. Fish gill parasite load has been shown to be positively associated with fish age, season, eutrophic water conditions [71], and temperature [72]. This may be explained by increased biofouling activity or biofilm formation over time on the gills or could be a response to parasite persistence. Unfortunately, we did not measure parasite abundances on the gill, but this would be an important area of research to examine the impact of parasite load on microbiome diversity, or vice versa. Understanding the effects of age on the microbiome was first demonstrated in African turquoise killifish where it was shown that microbiomes from older fish were associated with inflammation in the gut which could be rescued by fecal microbiome transplants from younger fish [73]. It has been suggested that during host ageing, gut communities of vertebrates may shift from commensal to pathogenic leading to increased inflammation and overall dysbiosis [74]. Our results indicate that microbial communities from other body sites may also be influenced by ageing or development of fish and is deserving of additional research. Additional host-associated explanatory variables not measured in our study, include diet or trophic level and host genotype. However, our assessment did determine at least based on mitochondria DNA, that the genetic population of mackerel was heterogeneous which further verifies the importance of environment and fish development stage on driving microbial communities.

Along with being most influenced by environmental and biological factors, overall the skin and gill communities were more similar phylogenetically to the sea water. Interestingly, of all mucosal sites, digesta samples were most similar to the sediment. This could be explained if the fish were feeding on benthic organisms [75] such as crustaceans buried in the sand. Although not quantified, we did find various types of crustaceans in the stomachs of the larger fish along with occasional gritty material which appeared to be sediment [76]. It is also possible that wave turbulence in nearshore environments where the fish were caught could also cause fish to be exposed to higher sediment levels through resuspension [77]. Since sediments are often repositories of decaying organic matter including anthropogenic contaminants, it is important to consider the negative health implications on a fish population as well as the potential human impacts associated with recreational fishing that occurs in near-shore locations, such as piers, and consumption of these fish.

Various potential pathogenic and beneficial microbes were persistently abundant across seasons which has important implications for climate change and aquaculture. *Photobacterium damselae* was present in skin and gut communities and in relatively high abundance compared to other microbes. High abundance relative to other microbes and prevalence across fish replicates could have important negative health implications as this is an important globally distributed [53] fish [78] pathogen causing bacterial septicemia that has also been implicated as an important genera for co-evolution in marine hosts [79]. If *P. damselae* is an important host-associated microbe, understanding the conditions by which it becomes pathogenic will be important in modeling fishery impacts. This pathogen has caused financial losses in marine fish farms across numerous species including yellowtail, gilthead seabream, and seabass [80, 81] and is thought to be transferred through water [82] to other species even infecting humans [83]. Chub mackerel are a very important forage fish consumed by many higher trophic level fish including tunas, billfish, and jacks which could have implications for trophic transfer of pathogens warranting future studies. Further, this microbe was most prevalent and abundant during the summer months suggesting it could be associated with high water temperatures and low nutrients. Extending this time series for another 3-10 years will be crucial to continue monitoring. While time series datasets exist for marine free-living microbial communities, few exist for marine vertebrates. Evaluating the extent by which exposure to marine pathogens influences disease is important for estimating impacts to fisheries. Further, as marine aquaculture activities continue to expand in coastal waters, farm monitoring through the host-microbiome could be an important tool for preventing disease outbreaks and massive losses. Experimental mesocosm studies could also be useful to model this marine vertebrate pathogen. Examples in other vertebrates of wide spread prevalence of opportunistic pathogens includes 20-80% carriage rates of *Staphylococcus aureus* in humans [84].

Some novel candidate symbiotic interactions were discovered when evaluating microbial ecology across the various mucosal sites. In the gills, several *Shewanella spp.* were highly prevalent in mackerel which is consistent with tropical fish microbiome studies [64] suggesting a potential symbiotic role. Some *Shewanella spp*. are common marine genera responsible for eicosapentaenoic acid (20:5n-3, an omega-3 polyunsaturated fatty acid) production [85, 86] and have been documented in freshwater fish [87]. Both Bacillus and Lactobacillus strains make up close to 50% of the microbial taxa in commercially available probiotics for aquaculture [88, 89]. Therefore, mucus from wild marine fish could provide a novel source of probiotics for use in the aquaculture industry.

In our study, we have for the first time evaluated the full mucosal microbiome of a single marine fish species. This design makes it the longest running wild marine fish microbiome study encompassing five body sites within a single species of fish caught over one year. The Pacific chub mackerel microbiome was primarily differentiated by mucosal body site. Environmental conditions and host biology primarily drives the skin and gill mucosal microbiomes with Chlorophyll a concentration, age, and temperature having the broadest effects. Our results provide the foundation to understanding natural microbiome variation of a wild marine fish which is economically important and provides a basis for asking how questions about how climate change may impact the marine fish microbiome in a positive or negative way.

## Supporting information

Supplemental data dictionary

## Acknowledgements

This work was supported by grants NSF (OCE-1313747) and NIEHS (P01-ES021921) to E.E.A. We thank the Center for Microbiome Innovation at UC San Diego for support through the Microbial Sciences Graduate Research Fellowship to J.J.M. We thank Karen Minich for graphic design assistance on Figure 1. We thank Shane Poplawski and Rachael Gominsky from JCVI for advice and assistance on using Oxford Nanopore sequencing.

