## Supplemental Figures for "Temporal, environmental, and biological drivers of the mucosal microbiome in a wild marine fish, *Scomber japonicus*"

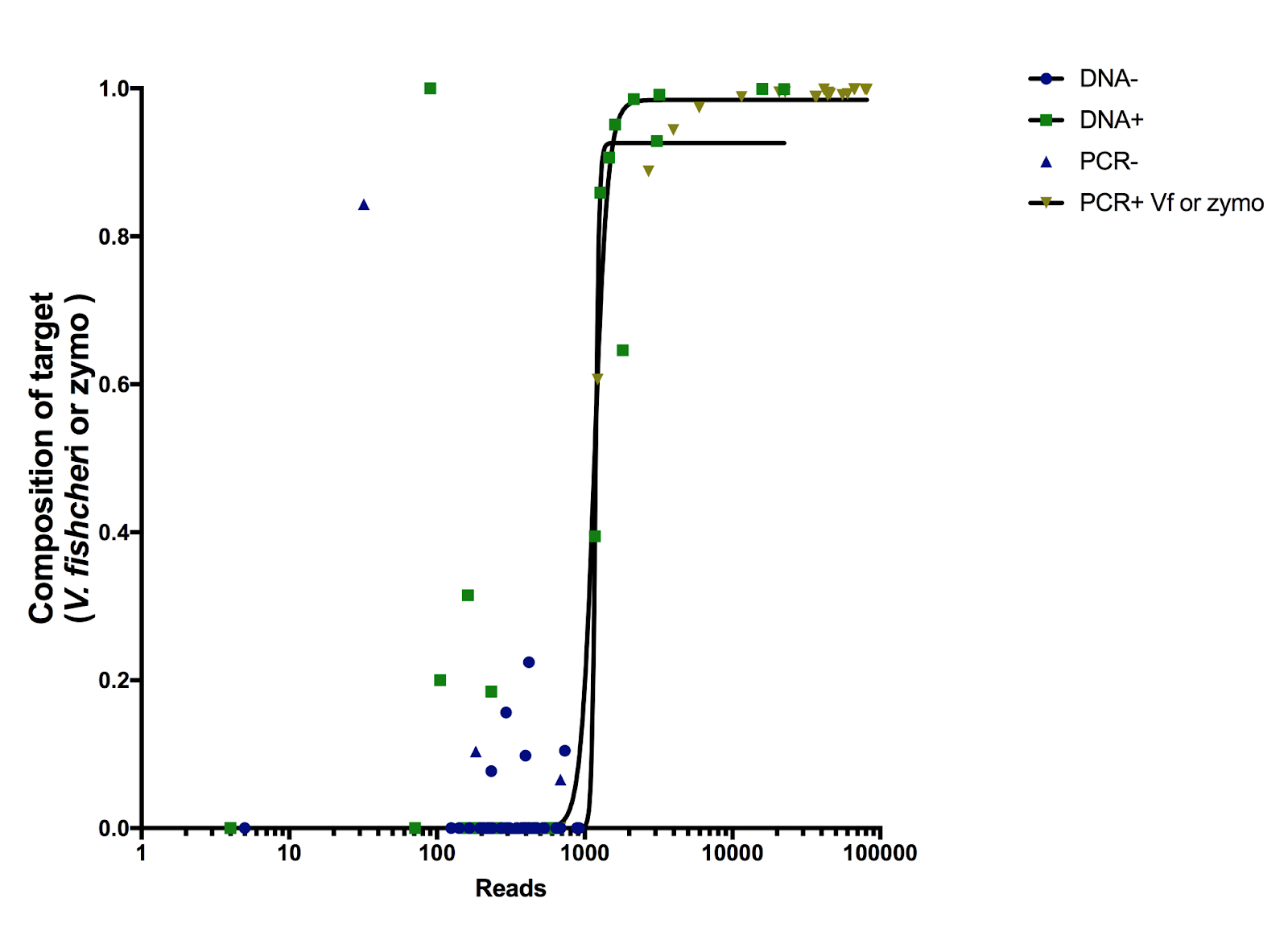

Supplementary Figure 1. Limit of detection titration curves generated from controls to determine sample exclusion criteria of 1362 reads.

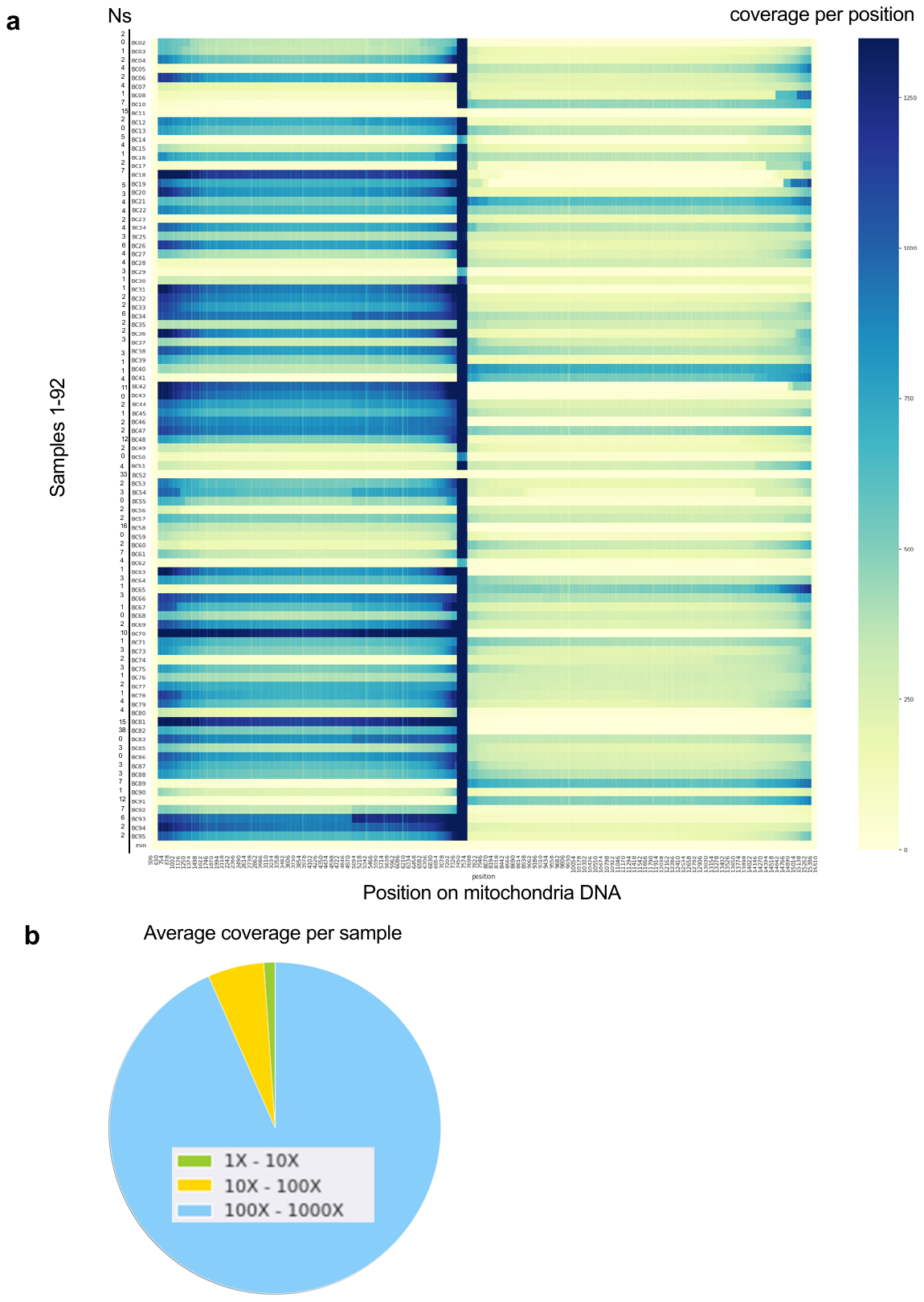

Supplementary Figure 2. Long read sequencing of mitochondria amplicon fragments 1 and 2. (a) Total coverage per base position across both fragments across 92 samples with number of Ns reported. (b) Summary statistic on coverage bins per samples.

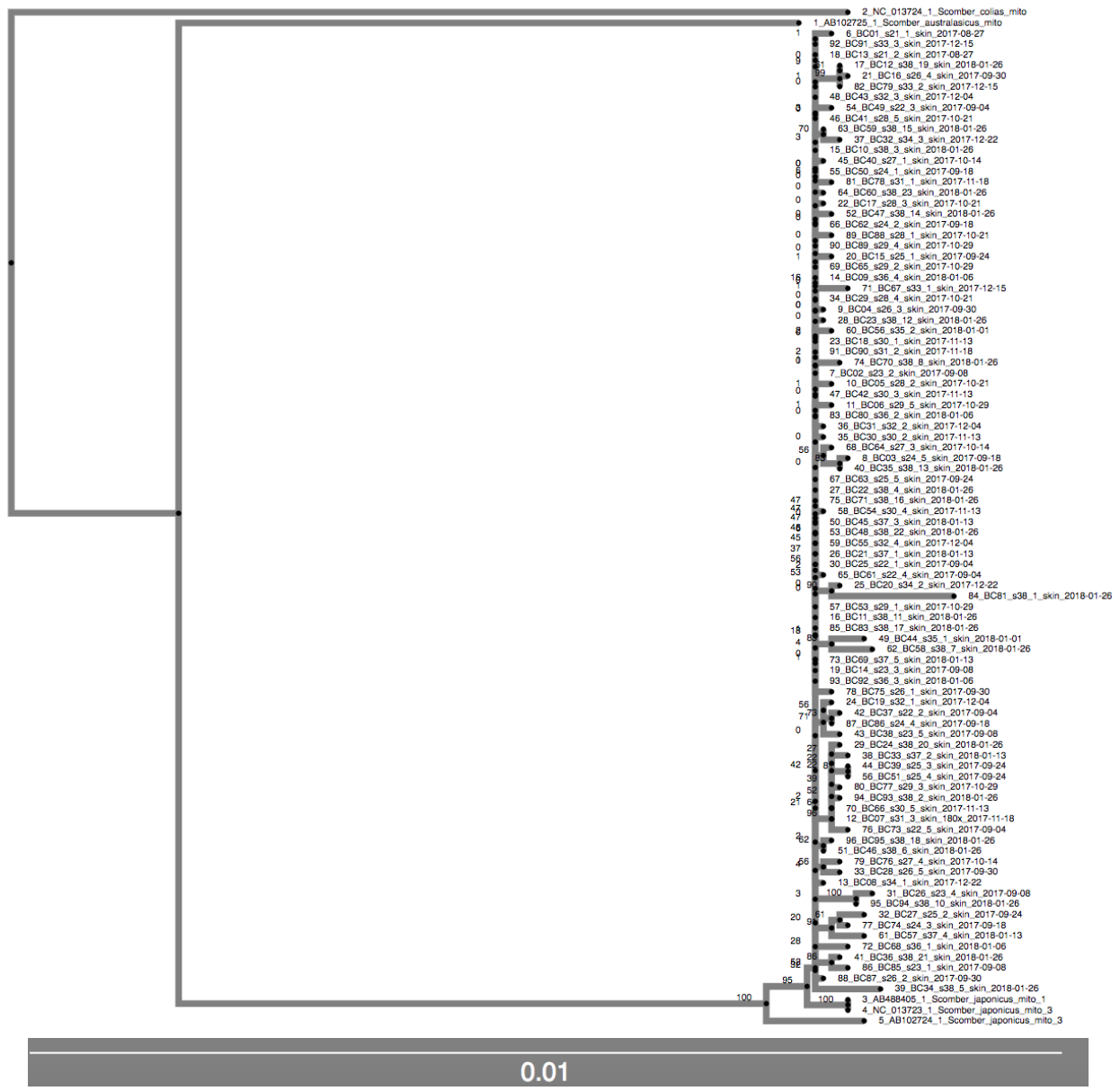

Supplementary Figure 3. Phylogenetic tree of near full length (14,769 bp) mitochondria DNA sequences across 91 *S. japonicus* sampled between Aug 27 2017 – Jan 26 2018. Outgroups include two other Scomber species: *S. colias* and *S. australasicus*.

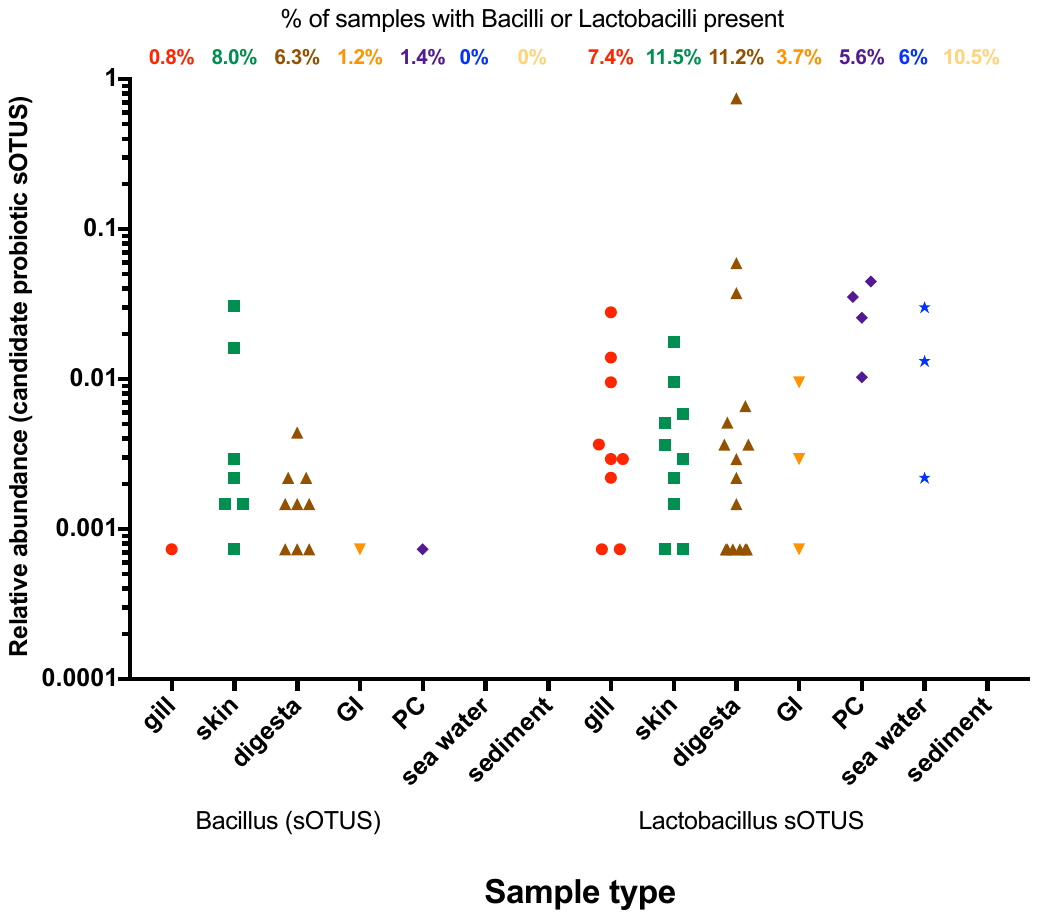

Supplementary Figure 4. Prevalence of candidate probiotics *Bacillus spp.* and *Lactobacillus spp*. on *S. japonicus* body sites throughout the sampling effort. Proportion of microbial community comprised of *Bacillus spp.* and *Lactobacillus spp.* sOTUs across the body sites and environment over the sampling effort of 1 year. Relative abundance is calculated as number of sOTU reads divided by 1362, the rarefaction number. Any samples with 0 *Bacillus* or *Lactobacillus* reads are considered under the detection limit and are not displayed.
